# Chandipura virus requires pro-survival RelA NF-κB function for its propagation

**DOI:** 10.1101/509893

**Authors:** Sachendra S. Bais, Yashika Ratra, Pramod K. Kushawaha, Soumen Basak

## Abstract

In response to infection by RNA viruses, mammalian cells typically activate RelA-containing NF-κB heterodimers, which induce genes encoding interferon-β and other antiviral mediators. Therefore, RelA is commonly thought to function as an anti-viral transcription factor. Notably, virus-specific mechanisms often modify mainstay immune pathways. Despite its human health relevance, how Chandipura virus (CHPV) per se interacts with the cellular signaling machinery has not been investigated. Here, we report that RelA deficiency abrogated antiviral gene expressions and yet surprisingly caused diminished growth of CHPV in mouse embryonic fibroblasts. Our experimental studies clarified that RelA-dependent synthesis of pro-survival factors restrained infection-inflicted cell death, and that exacerbated cell death processes prevented multiplication of CHPV in RelA-deficient cells. In sum, we identify a pro-viral function of the immune-activating transcription factor RelA NF-κB linked to its pro-survival properties.

**Highlights:**

- Lack of RelA NF-κB leads to reduced growth of CHPV ex vivo
- RelA deficiency exacerbates cell-death processes upon CHPV infection
- Inhibition of cell-death processes restores CHPV multiplication in RelA-deficient MEFs

## Introduction

Cytoplasmic RNA viruses continue to pose a threat to public health (Dye, 2014; Marston et al., 2014). In particular, Chandipura virus (CHPV) has been implicated in several recent outbreaks of acute encephalitis in India (Menghani et al., 2012). CHPV belongs to the Rhabdoviridae family and Vesiculovirus genus and possesses a single-stranded negative-sense RNA genome (Basak et al., 2007). Viral life cycle involves entry into host cells, the release of genetic material, synthesis of viral proteins as well as progeny genome RNA, assembly of mature virus particles and their exit from infected cells. CHPV also produces cytopathic effects and causes cell death. It was, in fact, suggested that neuronal death triggered by CHPV leads to neuropathogenesis in human patients (Ghosh et al., 2013).

Mammalian cells engage innate immune pathways for limiting cytoplasmic RNA virus infections (see Figure 1A). Previous biochemical studies involving vesicular stomatitis virus (VSV) and other prototype RNA viruses defined broadly the biochemical mechanism underlying antiviral host responses (Gerlier and Lyles, 2011; Scutigliani and Kikkert, 2017). In brief, viral nucleic acids through cognate receptors stimulate the activity of TANK-binding kinase 1 (TBK1) and IkappaB kinase ε (IKKε), which phosphorylate interferon regulatory factor 3 (IRF3) leading to its nuclear translocation. Moreover, viral infections activate the canonical nuclear factor-κB (NF-κB) pathway. In uninfected cells, RelA:p50 NF-κB heterodimers are sequestered in the cytoplasm by NF-κB inhibitor proteins (IκB) IκBα, IκBß and IκBε. Canonical signaling recruits a kinase complex consisting of NF-κB essential modulator (NEMO) and IKK2 (also known as IKKß), which phosphorylates IκBs. Consequent proteasomal degradation of IκBs liberates RelA:p50 into the nucleus. RelA:p50 and IRF3 cooperatively induce the expression of type-1 interferon (IFN) genes, particularly the one encoding IFNß. IFNß provides autocrine and paracrine signals through IFN-alpha/beta receptor (IFNAR) that upregulate hundreds of IFN-stimulated genes (ISGs) producing a robust antiviral cell state.

**Figure 1:**
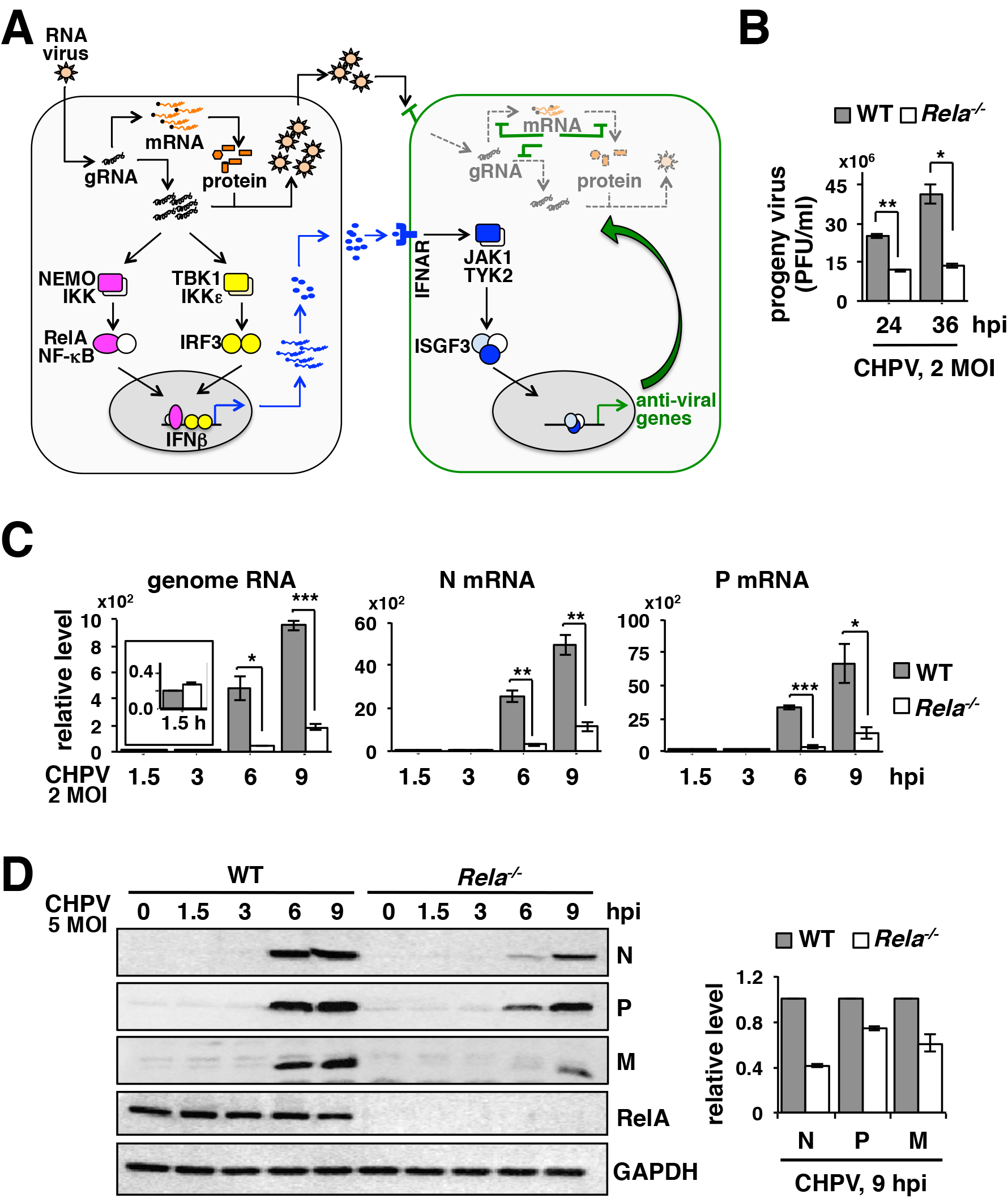
Genetically dissecting the role of RelA in regulating CHPV propagation. **A.** A current model for the cellular defense to RNA virus infections. In general, viral sensing activates the RelA NF-κB heterodimer, which in collaboration with IRF3 induces the expression of type 1 interferons, including IFNβ. Autocrine and paracrine signals of IFNβ through the cognate IFNAR restrict viral multiplication and produce an anti-viral state in neighbouring cells. Despite its relevance to human health, immune signaling pathways activated by CHPV *perse* have not been characterized. **B.** WT and *Rela^-/-^* MEFs were infected with CHPV at 2 MOI; culture supernatants were collected at the indicated times, and progeny virus yield was measured using plaque assay. Data represent means of three biological replicates ± SEM. **C.** RT-qPCR revealing the relative abundance of viral genomic RNA as well as N and P mRNAs in WT and *Rela^-/-^* MEFs infected with CHPV at 2 MOI for the indicated times. The abundance of viral RNAs was normalized to that of Actb mRNA. Data represent means of four biological replicates ± SEM. The inset shows the relative level of genome RNA subsequent to viral absorption in WT and RelA-deficient cells. **D.** WT and *Rela^-/-^* MEFs were infected with CHPV at 5 MOI, harvested at the indicated times post-infection and whole cell extracts were subjected to immunoblot analyses using antibodies against the indicated viral proteins. GAPDH served as a loading control. Right, densitometric analysis of the relative abundance of N, P and M proteins quantified from three independent experiments. The statistical significance was determined using two-tailed Student’s t-test. \**P* ≤ 0.05; \*\**P* ≤ 0.01; \*\*\**P*≤0.001

Genetic analyses causally linked the IRF pathway to cellular defence in relation to a number of cytoplasmic RNA viruses. For example, deficiency of *Irf3* diminished the production of IFNß by cells infected with VSV, Newcastle disease virus (NDV) or Dengue virus (DENV) (Chen et al., 2013; Sato et al., 2000). Similarly, knockout studies established a role of TBK1 and IKKε in phosphorylating IRF3 and activating the expression of IFNß in response to VSV, Sendai virus (SeV) and West Nile virus (WNV) (Benjamin et al., 2004; Fitzgerald et al., 2003; Fredericksen et al., 2004; Sharma et al., 2003). Consistently, a lack of IKKε or IRF3 potentiated the propagation of VSV, WNV or DENV ex vivo (Chen et al., 2013; Fredericksen et al., 2004; Sharma et al., 2003).

Canonical NF-κB signaling regulates a diverse range of cellular functions and mediates also the expression of pro-survival factors. Targeted deletion of gene encoding RelA in mice resulted in cellular apoptosis as well as necroptosis and caused embryonic lethality (Beg et al., 1995; Xu et al., 2018). Unavailability of adult mice lacking components of the canonical pathway impeded genetic dissection of NF-κB signaling in the context of virus infections. In a solitary study, Wang et al. (2010), infected mouse embryonic fibroblasts (MEFs) devoid of RelA with VSV or NDV at low multiplicity of infections (MOI) (Wang et al., 2010). Their study indicated that the canonical NF-κB pathway is important for early, but not late, expressions of interferon-β in infected cells and for suppressing the growth of these viruses. Of note, mainstay immune pathways are often modulated by additional virus-specific interventions (Taylor and Mossman, 2013). Despite its human health relevance, how CHPV per se interacts with the cellular signaling machinery has not been investigated.

Utilizing genetically tractable MEFs, here we examined the role of RelA in antiviral host responses to CHPV. CHPV induced nuclear translocation of RelA:p50 *via* the canonical NF-κB pathway. Indeed, RelA deficiency abrogated the expression of interferon-β in response to CHPV infection. Unexpectedly, infection of *Rela^-/-^* MEFs led to a decreased yield of progeny CHPV particles. Our experimental analyses clarified that this pro-viral NF-κB function was linked to the ability of RelA to suppress CHPV-induced cell death. In sum, we provide evidence that the pleiotropic transcription factor RelA may have pro-viral functions.

## Results and Discussion

### Diminished multiplication of CHPV in RelA-deficient cells

In general, mammalian cells engage NF-κB and IRF factors for inducing the expression of type 1 interferons, which mediate anti-viral functions (see Introduction and Figure 1A). Given RelA participates in diverse cellular processes, we asked if genetic deficiency of the pleiotropic factor RelA indeed led to an overall increase in the production of CHPV. To this end, we infected WT and *Rela^-/-^* MEFs with CHPV, and then measured the titre of progeny virus particles in the culture supernatant using the plaque assay (see Materials and Methods, and Figure S1A). Surprisingly, *Rela^-/-^* MEFs infected with CHPV at 2 MOI produced 2.2 fold less progeny virus particles at 24 hour post-infection (hpi) as compared to corresponding CHPV-infected WT cells; there was a further 3.1 fold reduction in the virus titre at 36hpi (Figure 1B). Next, we measured the abundance of viral genome RNA and mRNAs in infected cells using the reverse transcription-quantitative polymerase chain reaction (RT-qPCR). We found that CHPV genome RNA were abundant equivalently in WT and *Rela^-/-^* MEFs immediately after viral adsorption (inset in the left panel, Figure 1C). On-going viral replication in WT cells led to a robust increase in the abundance of CHPV genome RNA at 9hpi, whereas the increase was less obvious in *Rela^-/-^* MEFs (Figure 1C). Moreover, the abundance of mRNAs encoding viral nucleocapsid (N) protein as well as phosphoprotein (P) was markedly low in *Rela^-/-^* cells in relation to WT MEFs at both 6hpi and 9hpi. Our quantitative immunoblot analyses consistently showed a substantially diminished abundance of viral N, P and matrix (M) protein in *Rela^-/-^* MEFs at these time points (Figure 1D). Our study suggested that RelA promoted the production of progeny CHPV particles and positively impacted viral RNA synthesis ex vivo.

### Activation of canonical RelA NF-κB signaling in CHPV-infected cells

The NF-κB family consists of more than a dozen dimeric factors with RelA:p50 and RelB:p52 heterodimers being the most prevalent in majority of cell types (Mitchell et al., 2016). RelA:p50 and RelB:p52 are activated by the canonical and noncanonical NF-κB pathways, respectively. As such, RNA viruses trigger the canonical RelA:p50 activity, which along with IRF3 contributes to anti-viral IFN-β expressions. However, VSV also stimulates the noncanonical RelB:p52 activity, which actually inhibits the expression of IFN-β (Jin et al., 2014). Of note, RelB is encoded by a RelA-target gene (Basak et al., 2008). Indeed, RelA-deficiency diminishes basal expressions of RelB and impairs noncanonical RelB:p52 activation downstream of lymphotoxin-β receptor (Basak et al., 2008). We asked if CHPV engaged primarily the interferon-inhibitory noncanonical NF-κB pathway, whose weakening led to reduced CHPV yield in RelA-deficient cells.

We infected MEFs with CHPV at 2 MOI and then measured the nuclear NF-κB (NF-κBn) DNA binding activity using electrophoretic mobility shift assay (EMSA). WT MEFs elicited the NF-κBn activity within 3h of infection that was further augmented in a time course until 9h (Figure 2A). The onset of cytopathic effects dissuaded us from biochemically analyzing infected cells beyond the 9h time point. Our supershift analyses confirmed that this NF-κBn activity induced in WT MEFs was consisted of RelA:p50 (Figure 2B). Accordingly, *Rela^-/-^* as well as *Rela^-/-^Nfkb1^-/-^* cells, which lacked the expression of both RelA and p50, were unable to produce the NF-κBn activity upon CHPV infections (Figure 2C). Moreover, this CHPV-induced NF-κB activity temporally coincided with the NEMO-IKK activity and IκBα degradation in WT MEFs (Figure S2A and Figure S2B). Indeed, genetic deficiency of the canonical signal transducers NEMO or IKK2 abrogated completely NF-κBn induction upon CHPV infection (Figure 2D). Our immunoblot analyses revealed that CHPV infection stimulated IRF3 phosphorylation in WT MEFs and that RelA deficiency did not discernibly impact the virus-induced IRF3 activity (Figure 2E). Our analyses formally established that CHPV, akin to most other prototypical RNA viruses, engaged the canonical NF-κB pathway and triggered the RelA:p50 activity.

**Figure 2.**
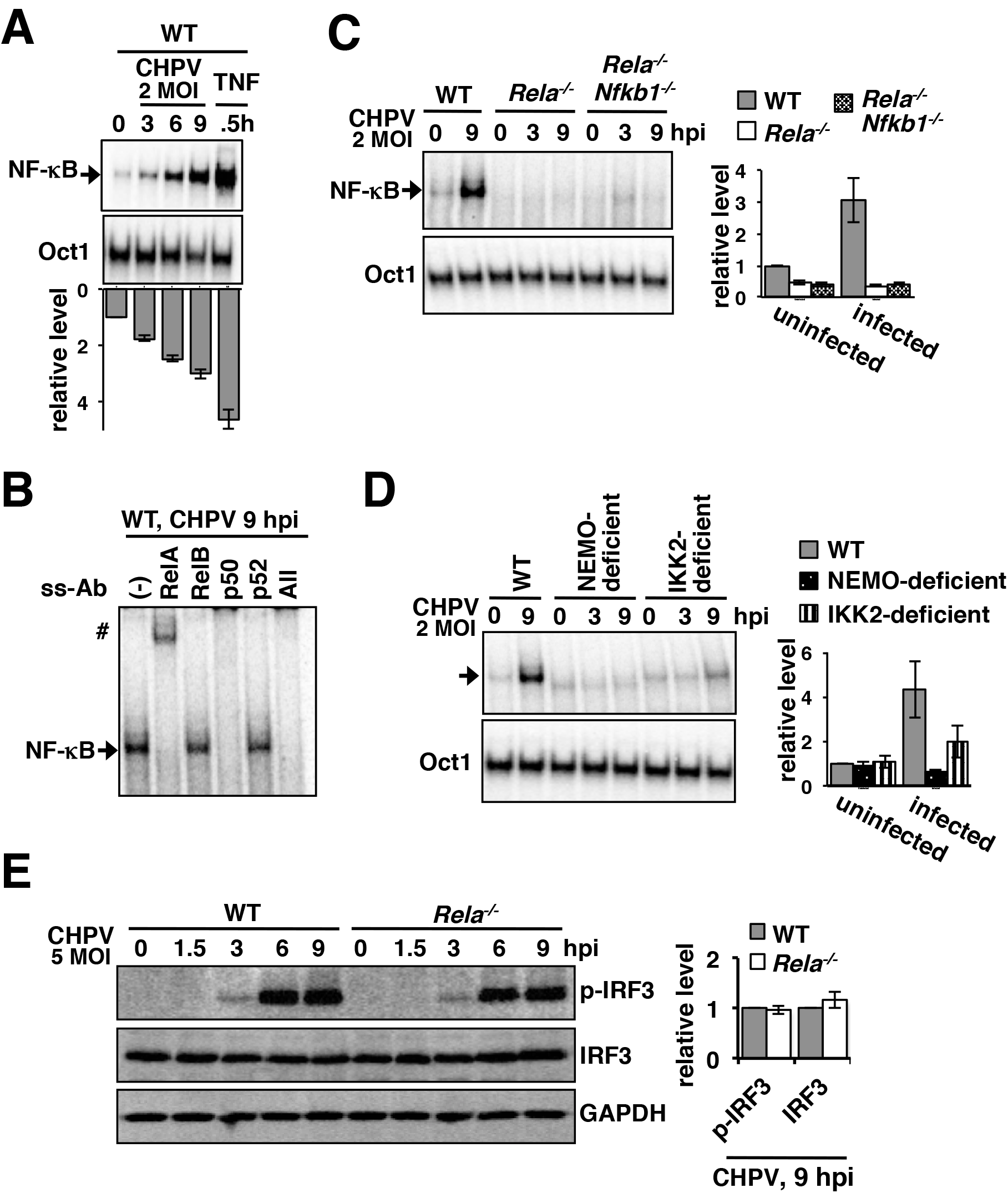
CHPV infection triggers the canonical RelA NF-κB signaling pathway. **A.** EMSA revealing the nuclear NF-κB activity (indicated by an arrow, top panel) induced in a time course in WT MEFs infected with CHPV at 2 MOI. Briefly, cells were harvested al the indicated times post-infection, nuclear extracts were prepared and examined for the presence of NF-κB DNA binding activity using a DNA probe containing kB site. Octl DNA binding served as loading control (bottom panel). The nuclear extract from TNF-treated cells was used as a positive control. The NF-κB activity from four independent experiments was quantified by densitometric analyses and presented in the bar plot as means ± SEM. **B.** Supershift analysis revealing the composition of the NF-κB dimers activated at 9h post-CHPV infection in WT MEFs. # represents supershifted bands. Anti-p50 antibody actually ablated the NF-κB DNA binding activity in EMSA. **C.and D.** NF-κB activities induced upon CHPV infection in WT MEFs and in various knockout MEFs devoid of one or more constituents of the canonical NF-κB pathway. For this study, immortalized MEFs were used. Right, the NF-κB activity from three independent experiments was quantified by densitometric analyses and presented in the accompanying bar plot as means ± SEM. **E.** Whole cell extracts derived from CHPV-infected (MOI 5) WT and *Rela^-/-^* MEFs were analysed for the abundance of p-IRF3 and IRF3 by immunoblotting. Right, the abundance of p-IRF3 and IRF3 proteins was quantified from three independent experiments and presented as means± SEM.

### Lack of RelA prevents anti-viral gene-expressions in response to CHPV infection

RelA activates genes with opposing biological functions (see https://www.bu.edu/nf-kb/ for a list of NF-κB-target genes). For example, RelA generally induces the expression of pro-inflammatory cytokines. However, RelA also activates the expression of antiinflammatory cytokines, such as interleukin-10, in specific cell types (Mosser and Zhang, 2008). Similarly, RelA stimulates the transcription of genes encoding important pro-survival factors, including Bcl2, cFLIP and cIAPs, in a wide variety of cells including cancerous cells. Curiously, RelA also upregulates the expression of FAS receptor and thereby sensitizing cells to specifically FasL-mediated apoptosis (Liu et al., 2012). It is thought that physiological and cellular context tunes NF-κB driven gene expressions. We asked if CHPV altered RelA-mediated controls of immune response genes, including that encoding interferons.

Our RT-qPCR analyses revealed a robust, three-log increase in the abundance of IFN-β mRNA in WT MEFs at 9h post-CHPV infection (Figure 3A), whereas RelA deficiency severely diminished IFN-β expressions. Autocrine and paracrine signals by type-1 interferons activate the expression of ISGs in virus-infected cells. Accordingly, CHPV infection led to gradual accumulation of mRNAs encoding ISG15 and OAS-1 in WT MEFs (Figure 3B). On the other hand, *Rela^-/-^* MEFs were defective for the expression of these genes. Virus-infected cells also produce pro-inflammatory chemokines and cytokines, which direct effector immune cells, including CD8 T cells and natural killer T cells, at the site of infection. Corroborating earlier studies showing a role of RelA in pro-inflammatory gene expressions, genes encoding CXCL10 and CXCL16 were induced in WT, but not in RelA-deficient, MEFs (Figure 3C). Furthermore, WT MEFs infected with CHPV upregulated mRNAs encoding various pro-survival factors, such as cIAP2, MnSOD and TRAF1; these genes were not activated in *Rela^-/-^* cells (Figure 3D). Of note, the abundance of cFOS mRNA, whose expression involves NF-κB-independent mechanisms, was not discernibly different between CHPV-infected WT and RelA-deficient MEFs (Figure 3E). Taken together, well-articulated transcriptional properties of RelA linked to its immune-activating and pro-survival functions were largely preserved in CHPV-infected cells.

**Figure 3:**
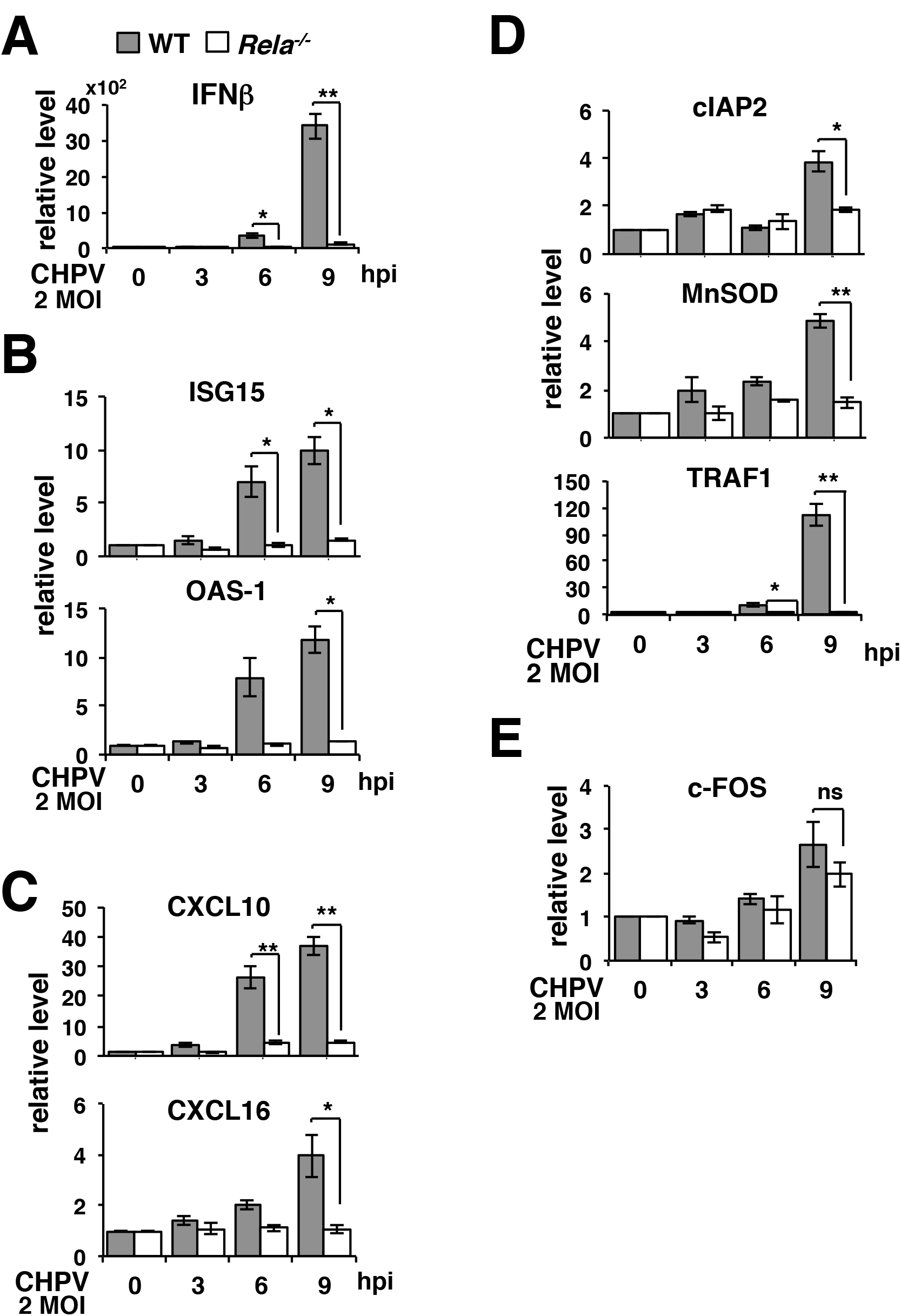
Investigating RelA NF-κB driven gene expressions in CHPV-infected cells. **A. to E.** RT-qPCR analyses revealing CHPV-induced expressions of mRNAs encoding IFNβ **(A)** or various ISGs **(B),** immune-activating chemokines **(C)** and pro-survival factors **(D)** in a time course in WT and *Rela^-/-^* MEFs infected at MOI 2. The expression of cFOS mRNA, which is encoded by an NF-κB-insensititve gene, was also scored **(E).** The abundance of mRNAs was normalized to that of Actb mRNA. Data represents means of three to four biological replicates ± SEM. \**P* ≤ 0.05; \*\**P* ≤ 0.01; \*\*\**P* ≤ 0.001 (paired two-tailed Student’s t-test).

### RelA-deficiency exacerbates cell-death processes in CHPV infected MEFs

Cell death abolishes the replicative niche of viruses and thereby serves as a host defense mechanism (Orzalli and Kagan, 2017). Cells infected with VSV and SeV activate the intrinsic apoptotic pathway where mitochondrial translocation of an IRF3-Bax complex causes cytochrome c release, caspase 9 activation and subsequent caspase 3 mediated cell death (Chattopadhyay et al., 2016). This cell death mechanism does not require transcriptionally active IRF3, and yet restrains effectively viral multiplication in vivo. RNA viruses also engage the extrinsic cell death pathway, which involves FAS-associated death domain (FADD)-mediated caspase 8 activation (Ghosh et al., 2013; Pearce and Lyles, 2009). In addition, recognition of influenza A virus (IAV) by cytosolic DNA-dependent activator of IRFs (DAI) stimulates receptor-interacting serine/threonine-protein kinase 3 (RIPK3) (Kesavardhana et al., 2017; Thapa et al., 2016). RIPK3 engages parallel cell death pathways - a kinase activity independent RIPK3 function reinforces extrinsic cell death signaling, while RIPK3 also phosphorylates mixed lineage kinase domain-like pseudokinase (MLKL), which promotes necroptosis. DAI-deficient mice fail to confine IAVs and succumb to infections (Thapa et al., 2016). VSV infections also trigger MLKL-dependent necroptosis (Wang et al., 2014). Viruses tend to counteract infection-induced cell death for promoting their growth. In our experiments, RelA upregulated pro-survival factors in CHPV-infected cells. We asked if RelA indeed suppressed cell death processes in CHPV-infected cells.

Our immunoblot analyses demonstrated that CHPV induced processing of procaspase 3 to caspase 3 in WT MEFs at 9hpi (Figure 4A). CHPV also promoted phosphorylation of MLKL at this time point. RelA deficiency not only accelerated the accumulation of caspase 3 and phosphorylation of MLKL in infected cells, but also augmented their abundance. CHPV did not alter the abundance of total MLKL, RIPK1, and RIPK3 in either WT or *Rela^-/-^* MEFs.

**Figure 4:**
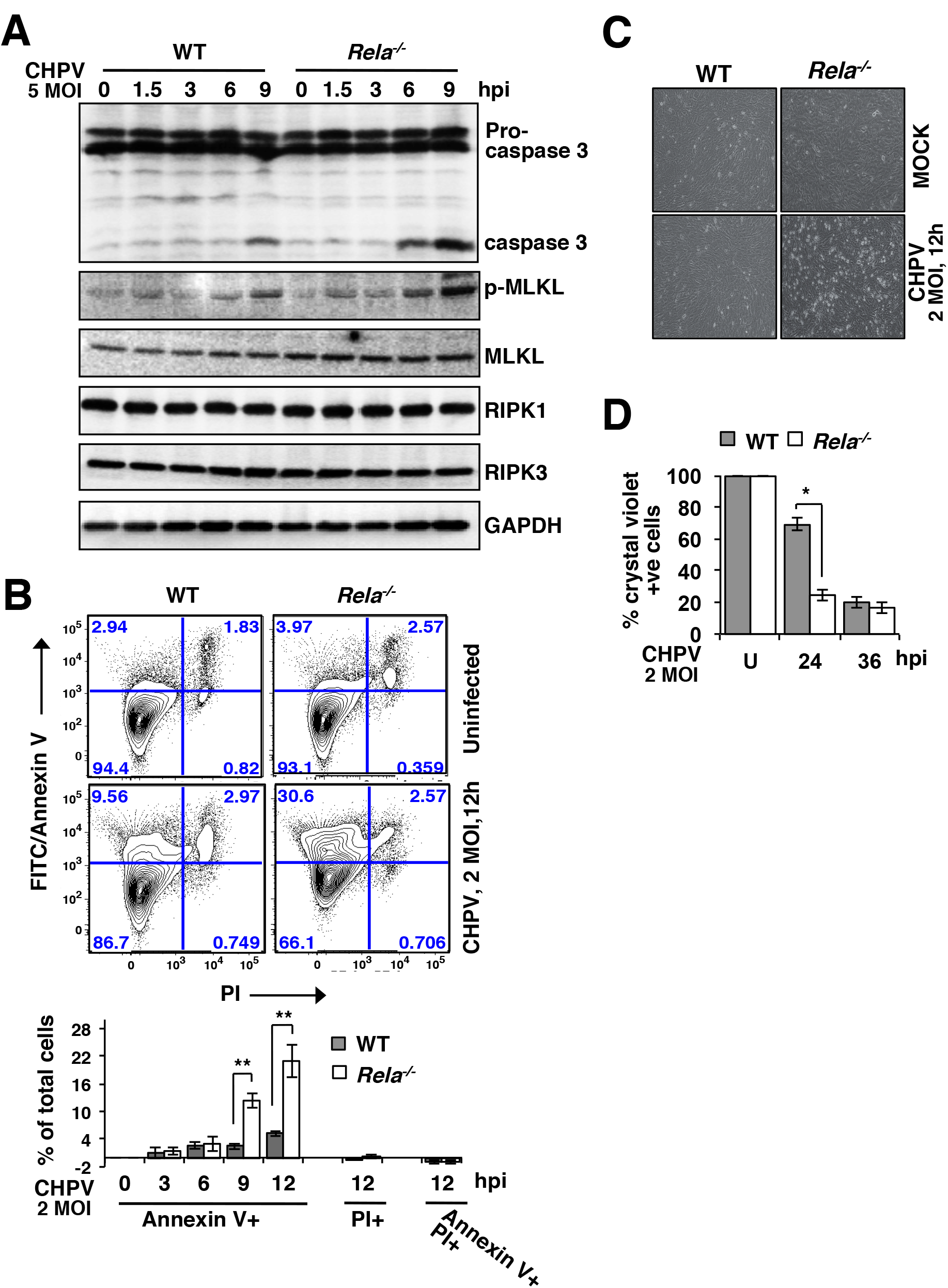
Charting cell-death processes in CHPV-infected WT and *Rela^-/-^* MEFs. **A.** Immunoblot analysis of whole cell extracts derived from WT and *Rela^-/-^* MEFs infected with CHPV at MOI 5 for the indicated times using antibodies against caspase 3, p-MLKL, MLKL, RIPK1, RIPK3 and GAPDH. **B.** CHPV-infected (MOI 2) WT and *Rela^-/-^* MEFs were stained with FITC/Annexin-V and PI before being subjected to FACS analysis. Uninfected cells were used as controls. Top, representative dot plots showing the prevalence of Annexin-V+, PI+, and Annexin-V+ PI+ cells in the uninfected and infected populations. Bottom, bar diagram revealing early cell-death quantified from three to six biological replicates, corrected for corresponding basal cell-death and presented as means± SEM. **C.** Representative light microscope images capturing CHPV-mediated cell rounding effects in WT and *Rela^-/-^*MEFs. **D.** Infection-induced cell-death at late time points were determined using crystal violet staining and presented relative to basal cell-death measured using uninfected MEFs. The plot represents average of three biological replicates ± SEM. \**P* ≤ 0.05; \*\**P* ≤ 0.01; \*\*\**P* ≤ 0.001 (paired two-tailed Student’s t-test).

Our fluorescence-activated cell sorting (FACS) analyses revealed that CHPV infection of WT MEFs for 12h only subtly increased the frequency of annexin V+ cells, which bear compromised membrane structure (Figure 4B). RelA deficiency produced ~12.0% annexin V + cells within 9h of infection and more than 20% annexin V+ cells at 12hpi. At these early time points, however, we were unable to detect either propidium iodide (PI)+ or annexin V+ PI+ cells that completely lack membrane integrity. Nonetheless, microscopic examination demonstrated discernible cytopathic effects at 12hpi in *Rela’*, but not WT, MEFs (Figure 4C). Crystal violet staining confirmed that CHPV triggered relatively rapid death of *Rela^-/-^* MEFs with less than 25% viable cells at 24hpi and ~15% live cells at 36hpi (Figure 4D). WT MEFs showed substantial delay in infection-induced death with close to 70% viable cells at 24hpi; however, prolonged infection reduced the cell viability to ~20% at 36hpi. Collectively, CHPV infection activated both apoptotic and necroptotic pathways that culminated into cell death and RelA suppressed these infection-induced cell death processes.

### Suppressing cell-death processes rescue CHPV multiplication in RelA-deficient MEFs

Next, we examined if suppressing apoptotic and necroptotic cell death processes could restore CHPV propagation in *Rela^-/-^* MEFs. To this end, we treated RelA-deficient cells for an hour with the pan-caspase inhibitor zVAD either alone or in combination with inhibitors of the RIPK3 kinase activity GSK843 or GSK872. Then these cells were infected with CHPV in the continuing presence of these inhibitors. Our immunoblot analyses revealed that zVAD treatment prevented the accumulation of mature caspase 3 in *Rela^-/-^* MEFs (Figure 5A). As described previously, inhibition of caspases enhanced the phosphorylation of MLKL in these cells (Nogusa et al., 2016). Use of RIPK3 kinase inhibitors along with zVAD abrogated both caspase activation and MLKL phosphorylation. Of note, the RIPK3 kinase pathway was shown to produce both annexin V+ and annexin V+ PI+ cells during the course of *Staphylococcus aureus* infection (Greenlee-Wacker et al., 2017). We found that zVAD alone was insufficient and a combinatorial treatment with zVAD and GSK843 was necessary for preventing the accumulation of annexin V+ cells undergoing death at 12h post-CHPV infection of RelA-deficient MEFs (Figure 5B and Figure S3). Notably, this combinatorial regime enhanced viral gene expressions - it augmented the abundance of genome RNA as well as N and P mRNAs at 9hpi (Figure 5C) - and CHPV propagation in *Rela^-/-^* MEFs (Figure 5D). Our analyses suggested broadly that caspases and MLKL cooperatively induce death of CHPV-infected cells, and that RelA promoted viral RNA syntheses and multiplication by restraining infection-inflicted cell death processes.

**Figure 5:**
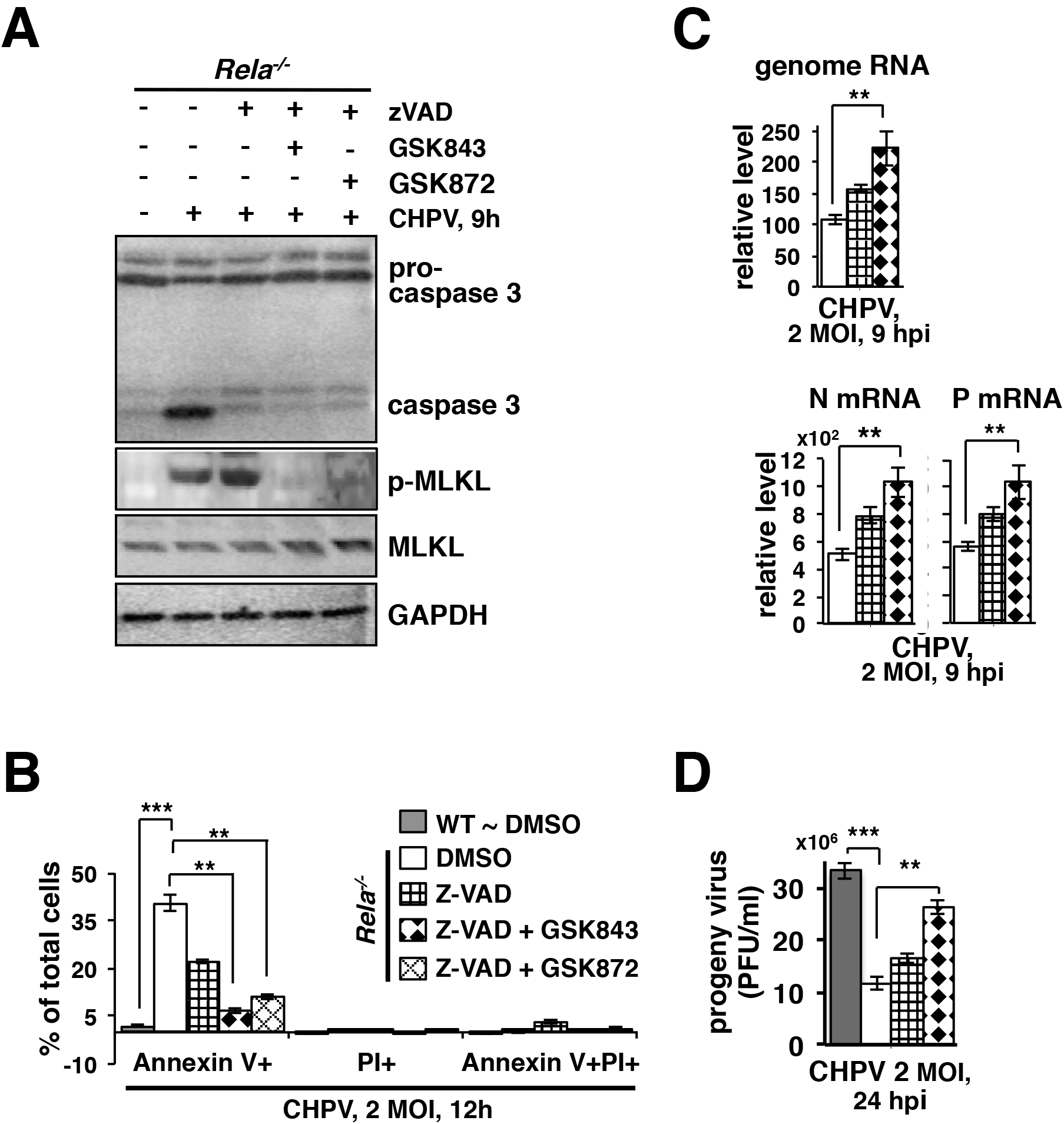
Investigating the effect of cell-death inhibitors on CHPV growth in *Rela^-/-^* MEFs. **A.** and **B.** WT and *Rela^-/-^* MEFs were infected in the presence of zVAD(OH)-FMK or in the concomitant presence of zVAD(OH)-FMK and GSK843 or zVAD(OH)-FMK and GSK872. Cells were harvested at 12hpi and subjected to immunoblot analyses using indicated antibodies **(A)** or examined for the presence of Annexin V+ cells using FACS **(B).** Bar diagram in **(B)** represents means of quantified data from four independent biological replicates ± SEM. **C. and D.** MEFs of indicated genotypes were infected with CHPV at 2 MOI in the absence or presence of cell-death inhibitors. The relative abundance of viral genomic RNA as well as N and P mRNA were determined using RT-qPCR **(C)**. Progeny virus yield in the culture supernatant at 24hpi was measured in the plaque assay **(D)**. The data represents the average of four **(C)** or three **(D)** biological replicates ± SEM. \**P* ≤ 0.05; \*\**P* ≤ 0.01; \*\*\**P* ≤ 0.001 (paired two-tailed Student’s t-test).

Type-1 interferons produced by infected cells restrict propagation of progeny virus particles into neighbouring cells. When used at high MOI, RNA viruses infect almost all the cells in a culture shortly after inoculation. We argue that high MOI used in our experiments rendered type-1 interferons irrelevant and in interferon-regulatory properties of RelA unimportant. In contrast, pro-survival RelA functions played a dominant role and supported the growth of cytopathic viruses in our studies.

Although CHPV infection activated caspase 3 and MLKL in *Rela^-/-^* MEFs within 6h (Figure 4A), it did not produce detectable PI+ dead cells at early time points (Figure 4B). However, transcription and replication of the CHPV genome were substantially attenuated in RelA-null cells even at 6hpi (Figure 1C). We propose that RelA potentiated CHPV growth involving two related mechanisms. For one, RelA safeguard the replicative niche of this intracellular pathogen by inhibiting cell death per se. RelA also modulated viral RNA syntheses presumably by protecting viral gene-expression machinery from the detrimental effects of caspases and MLKL. Indeed, cellular caspases were shown to cleave Crimean-Congo hemorrhagic fever virus nucleoproteins, which play an essential role in the viral replication (Karlberg et al., 2011). Despite the reduced abundance of CHPV N and P proteins in RelA-deficient cells, we were unable to detect fragmented viral proteins in our experiments. Future studies should further characterize how cell death mediators regulate CHPV gene expressions at early time points in an infection time course.

### RNA viruses and pleiotropic transcription factors

Cell death serves as an important anti-viral mechanism. Viruses have evolved mechanisms for preventing the death of infected cells, which serve as their replicative niche. DNA viruses typically encode pro-survival factors in their genome. For example, poxviruses express serine protease inhibitors serpins, which inhibit the activity of cellular caspases (Nichols et al., 2017). Cytomegaloviruses utilize virally-encoded inhibitors of Bax and Bak, which trigger the intrinsic apoptotic pathway, for preventing cell death (Brune and Andoniou, 2017). Additionally, MCMV-encoded M45 protein diminishes RIPK3-dependent necroptosis of infected cells (Upton et al., 2010). Oncogenic human herpesvirus 8, which is associated with Kaposi’s sarcoma, recruits a viral analogue of pro-survival factor cFLIP (Thurau et al., 2009). RNA viruses normally do not encode specialized pro-survival factors, presumably owing to relatively shorter genome length. Cells infected with viruses, including RNA viruses, elicit the RelA NF-κB activity, which upregulates the expression of anti-viral type-1 interferons. Our analyses revealed a rather counterintuitive pro-viral function of RelA particularly in the context of cellular infection with CHPV.

As such, RelA represents a pleiotropic transcription factor. In addition to activating the expression of IFN-ß, RelA also mediates the expression of genes encoding pro-survival factors. Indeed, RelA was shown to be capable of suppressing apoptotic as well as necroptotic cell death in response to a variety of death-inducing agents (Mitchell et al., 2016; Xu et al., 2018). Our investigation revealed that CHPV engaged both apoptotic and necroptotic pathways for inducing cell death and that RelA restrained death of CHPV-infected cells. We further identified that RelA-mediated suppression of infection-inflicted cell death augmented the yield of CHPV (Figure 6). Therefore, RNA viruses, which typically do not encode specialized pro-survival factors, appear to exploit pleiotropic cellular factors for extending the lifespan of infected cells. Our study also elucidated interesting contrast between two seemingly related transcription factors RelA and IRF3. IRF3 employs two distinct mechanisms for preventing viral propagation. It not only induces the expression of type-1 interferons in collaboration with RelA, but also triggers Bax-dependent cell death. However, RelA seems to play a dual role in the context of viral pathogenesis. By activating the expression of IFNß, RelA restricts viral growth; by inducing the expression of prosurvival factors, RelA amplifies the viral yield (Figure 6). We suggest that our finding has ramification for other similarly cytopathic RNA viruses and also in the development of antiviral therapeutic regimes.

**Figure 6:**
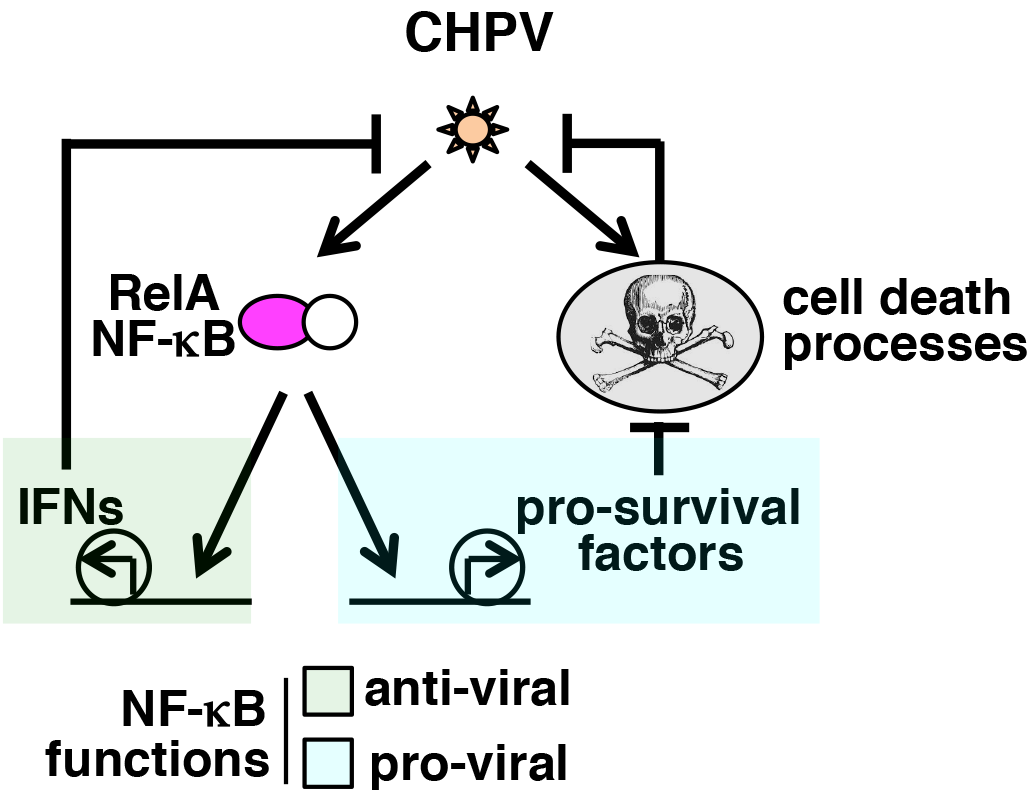
The proposed model explaining how RelA NF-κB promotes the multiplication of CHPV by restraining infection-induced cell-death.

## Materials and Methods

### Cells and viruses

WT and gene-deficient C57BL/6 mice were used in accordance with the guidelines of the Institutional Animal Ethics Committee of NII (approval no. #380/15). As described earlier (Roy et al., 2017), MEFs generated from E13.5 embryos were used either as primary cells or subsequent to their immortalization by the NIH-3T3 protocol. CHPV (strain 653514) was obtained from National Institute of Virology, Pune and propagated in VERO.

### Virus infection studies

Semi-confluent cultures of MEFs were incubated with CHPV for 1.5h in serum-free DMEM. Subsequent to viral adsorption, cells were washed with PBS and replenished with DMEM containing 10% BCS. At various times post-infection, either the culture supernatants were collected or the cells were harvested for further analyses.

### Viral plaque assay

Culture supernatants were collected from CHPV infected-MEFs at various times post-infection and analysed for the presence of progeny virus particles using the plaque assay. Briefly, monolayers of VERO E6 cells were infected with serial dilutions of the culture supernatants. After 60 min of incubation, inoculums were removed and cells were washed with PBS and overlaid with DMEM containing 1% low-melting agarose and 5% FCS. After another 24h of incubation, cells were fixed with formaldehyde and stained with crystal violet for visualizing viral plaques.

### Biochemical analyses

MEFs were harvested at different times post-infections; whole cell extracts and nuclear extracts were analysed by immunoblotting and EMSA, respectively (Roy et al., 2017). NF-κB-related antibodies have been described earlier (Roy et al., 2017). CHPV-related anti-N, anti-M and anti-P antibodies were obtained as a gift from NIV, Pune. Antibodies against p-IRF3 (#4947), IRF3 (#4302), Caspase 3 (#9665), p-MLKL (#62233), MLKL (#37705), RIPK3 (#95702), RIPK1 (#3493) and GAPDH (#2118) were from Cell Signaling Technologies, USA (MA, USA). The gel images were acquired using PhosphorImager (GE Amersham, UK) and quantified using ImageQuant 5.2. The IKK2 kinase assay was performed as described previously (Banoth et al., 2015).

### RNA analyses

Total RNA was isolated from infected MEFs and RT-qPCR analyses was conducted, as described (Roy et al., 2017). Supplementary table 1 provides a detailed description of primers used in this study for determining the abundance of CHPV genomes and CHPV-encoded as well as host-derived mRNAs in virus-infected cells.

### Cell death studies

Infection-inflicted early cell death was quantified by FACS analyses subsequent to staining of MEFs with FITC/Annexin-V and PI using apoptosis detection kit from BD Biosciences (# 556547). Data was obtained using VERSE (BD Bioscience, NJ, USA) and analyzed in FlowJo 9.8.3. Infection-induced cell-death at late time points were determined by crystal violet staining, as described earlier (Zong et al., 1998). In certain experiments, cells were incubated with various cell death inhibitors, including zVAD (OH)-FMK ((#14467, Cayman chemicals, MI, USA), GSK843 (#4898, Aobious, MA, USA) and GSK872 (#2673, Biovision, CA, USA), for 1hr before virus infections.

### Statistical analysis

Error bars were shown as S.E.M. of three or more experimental replicates. Quantified data are means ± SEM, and two-tailed Student’s t-test was used for verifying statistical significance.

## Supplemental information

Supplemental Information includes three figures and a table associated with the main text.

## Author contributions

SSB carried out the wet laboratory experiments with the help from YR, and PKK and supervision from SB. SB conceived and supervised the overall research. SSB wrote the manuscript with SB. The authors declare no conflict of interest.

## Acknowledgments

We thank Prof. D Chattopadhyay, Amity University, Kolkata for insightful discussions and help with reagents. We thank V. Kumar SIL, NII for technical help and P. Nagarajan from SAF, NII for help with animal husbandry. Virus research in PI’s laboratory is funded by NII-core. SSB and YR, thank CSIR and DBT, respectively, for research fellowships.

