## Supplemental information for "Chandipura virus requires pro-survival RelA NF-κB function for its propagation"

### Supplementary Material

Figure S1: Investigating the propagation of RNA viruses in MEFs

Figure S2: CHPV-activated canonical NF-κB signaling in MEFs

Figure S3: Preventing CHPV-mediated cell death using various inhibitors

Table S1. List of the primers used in our quantitative real-time PCR

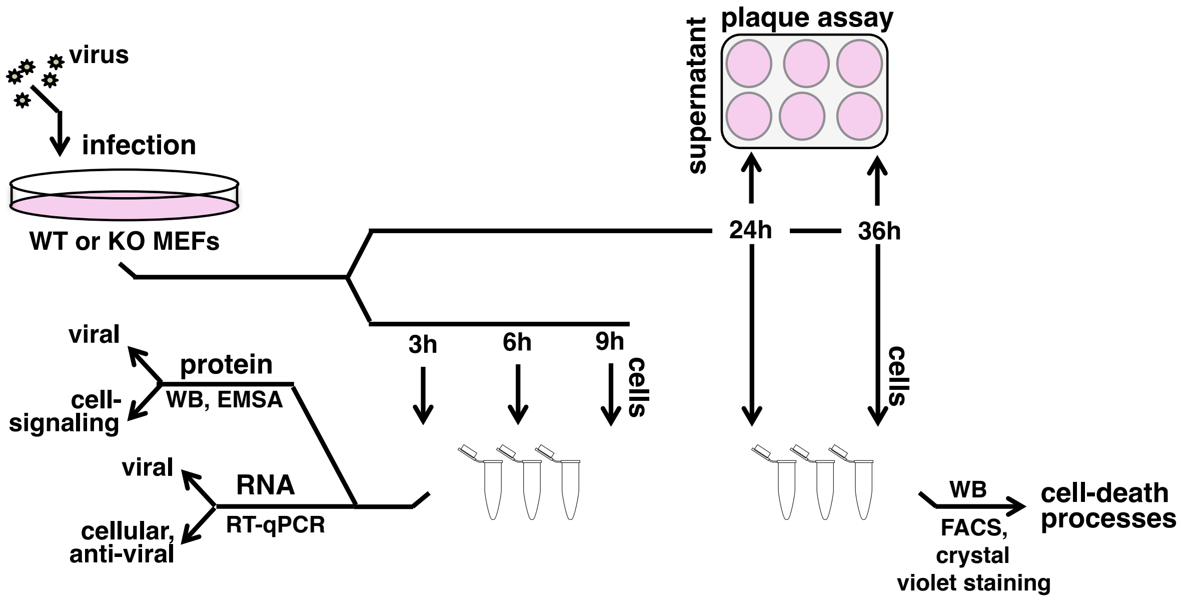


**Figure S1: Investigating the propagation of RNA viruses in MEFs.**

Schematic presentation of viral infection studies in MEFs. WT and knockout cells were infected with CHPV at different MOIs. Infected MEFs were harvested at 3h, 6h, and 9h post-infection and subjected to protein and RNA analyses. Alternately, cells were harvested at 12h or 24h or 36h post-infection and analyzed for cell-death process, and corresponding culture supernatants were tested for the presence of viral particles using the plaque assay.

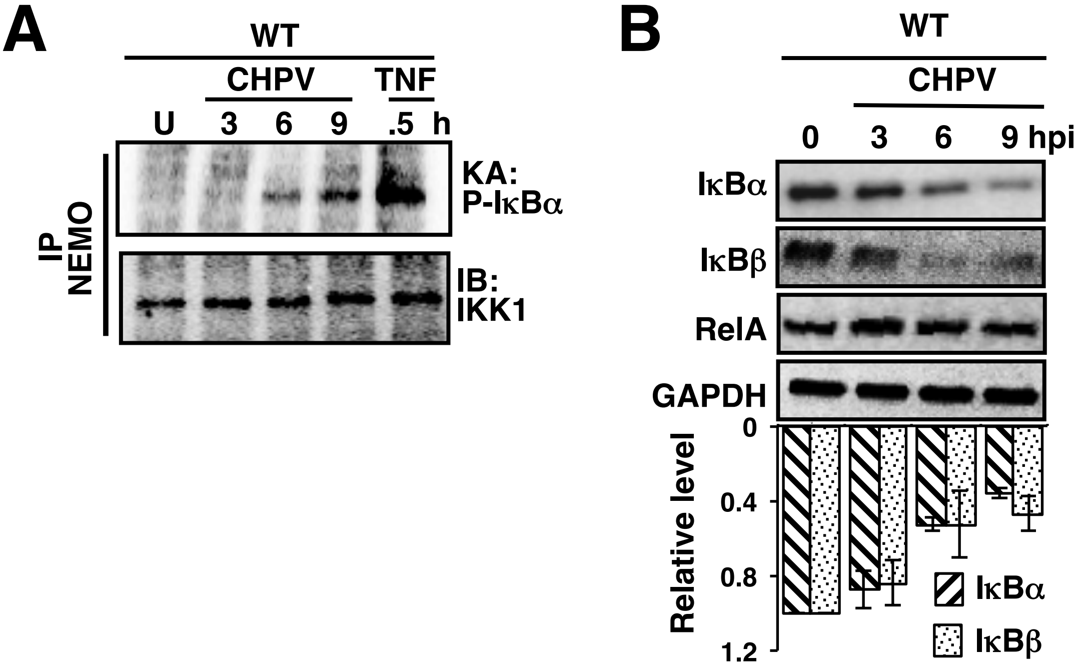


**Figure S2: CHPV-activated canonical NF-κB signaling in MEFs.**

**A.** Time course analyses revealing the NEMO-IKK2 activity induced upon CHPV infection at MOI 5. Briefly, NEMO co-immunoprecipitates derived from CHPV-infected MEFs were incubated with recombinant GST-IκBα and γ-32P ATP; the resultant mixture were resolved onto SDS-PAGE. Co immunoprecipitated IKK1 served as a loading control.

**B.** WT and *Rela*$^{-/-}$ MEFs were infected with CHPV at 5 MOI, harvested at the indicated times post-infection and whole cell extracts were subjected to immunoblot analyses using antibodies against RelA, IκBα and IκBβ. GAPDH served as a loading control. Bottom, densitometric analysis of the relative abundance of IκBα and IκBβ quantified from three independent experiments.

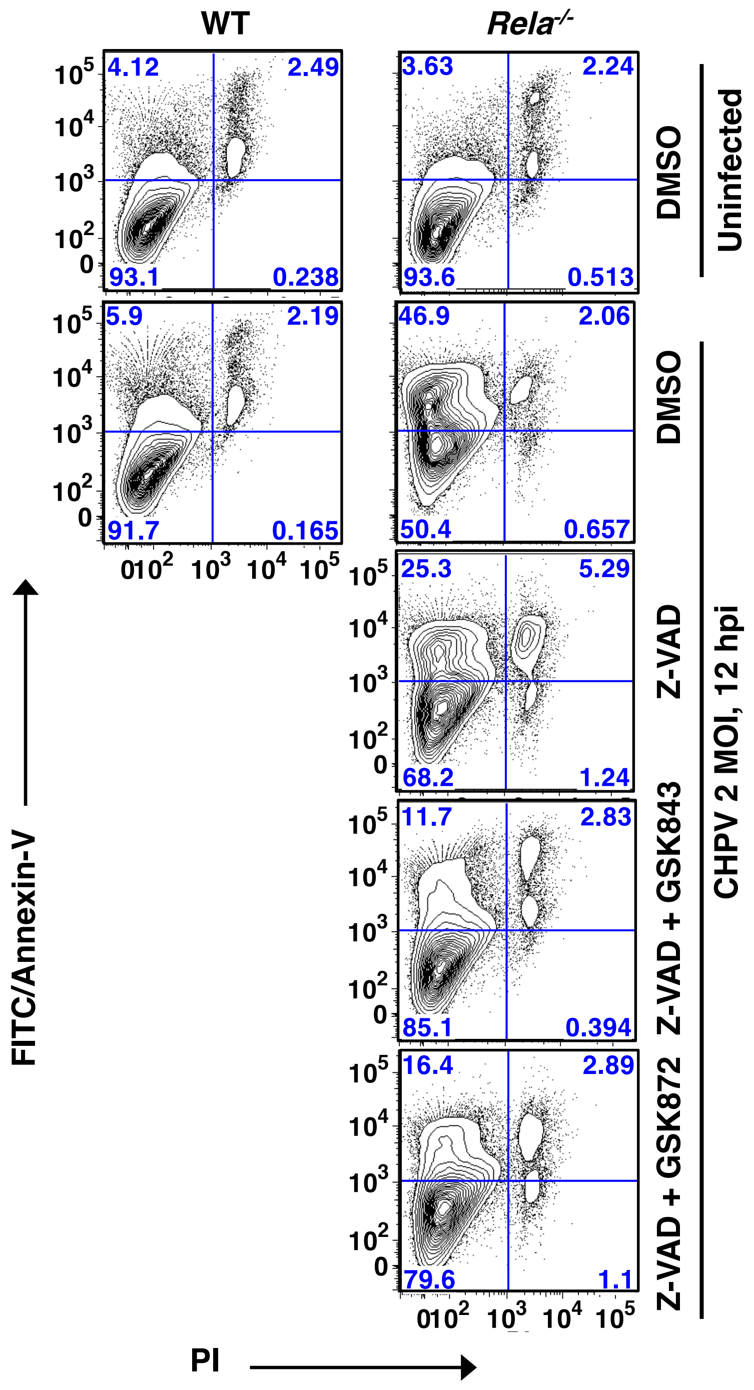


**Figure S3: Preventing CHPV-mediated cell death using various inhibitors.**

WT and *Rela*$^{-/-}$ MEFs were infected with CHPV in the presence of Z-VAD(OH)-FMK or in the concomitant presence of Z-VAD(OH)-FMK and GSK843 or Z-VAD(OH)-FMK and GSK872. Cells were harvested at 12 hpi and examined for the presence of Annexin V+ cells or PI+ cells or both using FACS.

**Table S1. List of the primers used in our quantitative real-time PCR.**

| Gene Name | Forward 5'→3' | Reverse 5'→3' |
|---|---|---|
| **CHPV Genome** | CGAGTGAACTCAGTTGCAGAG | GAATCGAGAGTGTCCTGAAGC |
| **CHPV-N** | GATTTGTTGCGGATGATGAC | CCAGAAATGGAAACTGGGAT |
| **CHPV-P** | CTCTCCGTCTGATCCACCTT | TCAATCCAGCAATGACCAGT |
| **mIFNβ** | CCGGACTTCAAGATCCCTATGGA | TGGCAAAGGCAGTGTAACTCTTC |
| **ISG-15** | AGCTCCATGTCGGTGTCAG | GAAGGTCAGCCAGAACAGGT |
| **OAS-1** | AGGGGCATTTGCTGCTCTGC | GGGCACCTGCTGTGGTTTATTG |
| **CXCL-10** | AGACATCCCGAGCCAACCTT | GTTAAGGAGCCCTTTTAGAC |
| **CXCL-16** | TGGAACTGGTCATGGGAAGAG | GGGTACTGGCTTGAGGCAAA |
| **cIAP-2** | GAAGTGGGCTGCGGTATCA | GCGCTGTCTTGAACCATGTTC |
| **MnSOD** | TCAATGGTGGGGGACATATT | GCTTGATAGCCTCCAGCAAC |
| **TRAF1** | GGAGGCATCCTTTGATGGTA | AGGGACAGGTGGGTCTTCTT |
| **cFOS** | CCTTCGGATTCTCCGTTTCTCT | TGGTGAAGACCGTGTCAGGA |
| **Actin** | CCAACCGTGAAAAGATGAC | GTACGACCAGAGGCATACAG |
